# Dehydration and infection elicit increased feeding in the western flower thrips, *Frankliniella occidentalis*, likely triggered by glycogen depletion

**DOI:** 10.1101/2022.07.14.499040

**Authors:** Samuel T. Bailey, Alekhya Kondragunta, Hyojin A. Choi, Jinlong Han, Dorith Rotenberg, Diane E. Ullman, Joshua B. Benoit

## Abstract

We examined water balance characteristics and influence of desiccating conditions on adult western flower thrips (*Frankliniella occidentalis*) physiology and behavior. Western flower thrips are globally invasive and likely to contend with shifts in water availability across their expansive geographic range. Basic water balance characteristics, including water mass and dry mass, were established for adult males and females, revealing a distinct sexual dimorphism wherein females are larger, but males retain a larger percentage of their mass as body water. Males lose relative water mass more quickly and their survival times are shorter when compared to females. RNA-seq analysis identified significant enrichment of factors associated with carbohydrate transport and metabolism in dehydrated males and females. A reduction of glycogen reserves was confirmed during dehydration. The probability of thrips feeding significantly increased when desiccation was a factor. Lastly, infection with *Tomato spotted wilt orthotospovirus* (TSWV), a principal plant-pathogenic virus transmitted by *F. occidentalis*, did not have a consistent and apparent influence on desiccation tolerance; however, a reduction in glycogen reserves, and an increase in feeding activity in infected thrips, very similar to that observed in dehydrated thrips, was observed. Our results establish the fundamental water balance characteristics of adult thrips, and indicate that dehydration significantly influences the survivorship and feeding behavior of thrips; crucial factors that contribute to their capacity to spread disease.

## Introduction

Among over five thousand known species of thrips worldwide, only a few species of thrips serve as major crop pests. Western flower thrips, *Frankliniella occidentalis* (Pergande) are significant, globally invasive, generalist pests of agriculture (Lewis 1997, Kindt et al. 2003). *Frankliniella occidentalis* is detrimental to a multitude of crops, as well as uncultivated plants spanning over 250 species, including fruiting and leafy vegetables, ornamentals, tree and berry fruits, and cotton (Reitz 2009, Nyasani et al. 2013). Their large, diverse host range, high fecundity, arrhenotokous parthenogenic reproduction, short generation time, and preference for concealed spaces, make thrips highly effective pests (He et al. 2020). Thrips cause direct damage to plants by piercing and sucking out the contents of plant cells (Childers 1997). Thrips feeding can cause stunting of plant growth or fruit scarring leading to crop loss, which can quickly become unmanageable, especially when aesthetic damage to the crop is not acceptable (Welter et al. 1990, Childers 1997, Reitz et al. 2020). Further injury is endured during oviposition because thrips insert their eggs into plant tissue using their saw-like ovipositor (Whittaker and Lewis 1975, Childers 1997).

Thrips transmit a myriad of plant-infecting orthotospoviruses through salivation while feeding, much like most other insect vectors (Rotenberg et al. 2015). The most economically significant virus spread by thrips, *tomato spotted wilt orthotospovirus* (TSWV), caused an estimated $1.4 billion in losses in the U.S. over a 10 year period and an annual worldwide loss of over $1 billion (Rybicki 2014, Riley et al. 2011). TSWV is acquired by thrips during the larval stage and is persistently transmitted in a circulative-propagative manner (Ullman et al. 1997); however, only thrips that acquire this virus as larvae are able to transmit as adults. Infection of the principal salivary glands is a prerequisite for the insect to inoculate plants with the virus, and this does not occur until very late in the second larval instar, just prior to pupation (Montero-Astúa, et al. 2016). The principal salivary glands do not become infected in adults that did not acquire the virus in the larval stages. There are multiple hypotheses that offer an explanation as to why the virus can only disseminate to the principal salivary glands during the larval stage, some suggesting differences in morphology that bring the salivary glands in proximity to virus-infected organs, and others proposing that receptor mediation, or other physiological differences may play a role in this phenomenon (Moritz et al. 2004, Schneweis et al. 2017, Ullman et al. 1992). In addition to transmitting many economically important plant pathogens, thrips are also increasingly resistant to commonly used pesticides, creating a need for novel control methods (Morse and Hoddle 2006). Despite their status as a global pest, little is known about their physiological stress limits.

The ability to tolerate environmental stress is vital in order for a species to establish in a new locality (Zerebecki et al. 2011). The native geographic range of *F. occidentalis* is western North America, which was likely maintained by the vast grasslands of the central US by harsh winters and the predominance of grasses and scarcity of flowers in the summer (Kirk and Terry 2003). Their distribution as a globally invasive crop pest likely began during increased international trade of horticultural materials in the 1960s (Kirk and Terry 2003). Increasing mean temperatures in the US could have also facilitated their spread (Horton 1995), although most evidence points to human activity (Mound 1983). Thrips population genetics in China revealed that they were introduced multiple times, supporting the idea that human activities are a primary driver in their global spread (Cao et al. 2017).

Water availability and temperature are the two most important abiotic factors that influence the abundance and distribution of insects in a given environment (Chown and Nicholson 2004). Due to their inability to survive harsh winters (McDonald et al. 1998), the western flower thrips likely needed the refuge provided by glasshouses in order to expand into new locales that they otherwise would not have been able to. This species is not obligately diapasual, providing further support that colder climates represent a physical barrier that restricts the spread (McDonald et al. 1998), although rapid cold hardening against low temperatures has been documented in thrips (McDonald et al. 1997). Due to their small size and thigmotactic nature, the use of favorable microclimates might provide them some capability for overwintering (Kirk and Terry 2003). The more mild winters of the western US and use of glasshouses as a refuge from fluctuating climates likely allowed the western flower thrips to colonize areas with colder climates. Rising global temperatures may facilitate an expansion of the geographic range of thrips.

When migrating between plant hosts and expanding their range, aside from temperature, thrips are at risk for dehydration due to their minute body size and consequently high surface area to volume ratio (Wharton 1985, Chown and Nicholson 2004). Despite this, the basic water balance characteristics and preventative measures used by thrips to resist dehydration and prevent damage from dehydration are currently unknown. In general, water balance is defined simply as the relationship between water gain and water loss (Wharton 1985). Water is gained by ingestion, metabolism, and, in some arthropods, absorption of water vapor (O’Donnel and Machin 1988). Loss occurs passively through the cuticle and through processes such as respiration, defecation, and secretion (Wharton 1985, Hadley 1994). In arthropods, water balance retention and damage reduction has been documented to be facilitated in a multitude of ways, such as behavioral adaptations, reduction of metabolic rate, increased sclerotization of the cuticle, alteration of cuticular hydrocarbons, discontinuous gas exchange, upregulated expression of heat shock proteins, and increased stores of protective osmolytes (Benoit et al. 2005, Benoit 2009, Bursell 1957, Hadley 1981). Maintaining water balance through water uptake is a key attribute that increases feeding in other disease vectoring arthropods, such as mosquitoes (Hagan et al. 2018). Such behavioral shifts in response to dehydration could cause an increase in crop yield damage by higher rates of feeding and viral spread during and following dry periods. Additionally, the plant stress hypothesis proposes that drought can increase herbivore fitness and abundance by increasing the nutritional quality of plants or reducing plant defenses, depending on the severity of the drought (Dale et al. 2017). In particular, it is postulated that generalist pests will succeed more in severe drought and specialist pests will thrive in moderate droughts. There is strong support for carbon and nitrogen compounds in plant tissues fluctuating in response to drought conditions (increasing during moderate droughts and decreasing during severe droughts) (Farooq et al. 2009). This implies that the nutritional quality of the plant likely increases during moderate drought, so insect species that can tolerate the higher level of defense compounds, mainly specialist feeders, will thrive. In more severe drought, generalists will thrive because the plant will have weakened defenses. Due to this, as temperatures increase due to climate change and droughts increase in frequency, it is hypothesized that generalist pests, like the western flower thrips, will thrive (Gely et al. 2020). As insects that spend most of their lives on a host plant, thrips likely obtain most, if not all, of their water through feeding, thus water balance should be accounted for when assessing feeding, vectorial capacity, and disease transmission (Holmes and Benoit 2019, Holmes et al. 2022).

Many plant pathogens, including plant viruses, are transmitted by insects and are solely reliant on these vectors to facilitate their spread. Consequently, it is hypothetically beneficial for these viruses to modulate aspects of the vector’s behavior and/or physiology as a means of self-preservation (Nachappa et al. 2020). In the case of TSWV, which replicates in both the thrips vector and plant host, modulation can occur in either host, with the interaction between hosts ultimately being beneficial to the virus (Reitz et al. 2020). Specifically, infection could affect the fitness of its vectors directly or infection of the plant may alter its suitability as a host for thrips (Stumpf and Kennedy 2007). TSWV infection has been shown to directly benefit thrips by increasing survival, as well as increasing the number and duration of feeding probes by infected adult males (Stumpf and Kennedy 2007, Belliure et al. 2004, Shrestha et al., 2012, Ogada et al., 2012, Stafford et al. 2011). Indirect plant-mediated effects include benefits to thrips such as increased development, fecundity, and population growth, as well as reducing host-plant defenses against thrips (Stumpf and Kennedy 2007, Belliure et al. 2004, Carter 1939, Shrestha et al., 2012, Nachappa et al. 2020). Viral infection may also modulate insect thermal tolerance, which is seen in other insects (Porras et al. 2020, Xu et al. 2016, Pusag et al. 2012). Numerous studies support that there is cross tolerance between dehydration and cold (Sinclair et al. 2007, Sinclair et al. 2013). Due to the link between thermal and dehydration stress, dehydration tolerance may also be influenced similarly to thermal tolerance due to viral infection. Although there are studies that examined the dynamics between viral infection and plant water stress (Nachappa et al. 2016), no studies have directly examined the dynamics between viral infection and dehydration in insects and how this might impact other phenotypes governing feeding and virus transmission.

In this study, we examined the water balance characteristics of adult western flower thrips in relation to infection with TSWV. As a globally invasive pest, it is critical to understand the dehydration tolerance of thrips, because dehydration is a significant abiotic stressor that impacts geographical distribution, reproductive capacity, longevity, growth rates, and feeding behavior (Chown et al. 2011, Albers and Bradley 2006, Finch et al. 2020, Benoit et al. 2010, Hera and Reichert 2021, Hagan et al. 2018). Various viruses are known to influence the physiology and behavior of insects; thus, surveying the potential interplay between dehydration tolerance and TSWV infection may reveal a critical host-virus interaction leading to increased insect vector fitness and/or likelihood to spread disease. To examine water balance requirements and the influence of TSWV on these characteristics, we determined water content, water loss rates, and survival, as well as the effect of dehydration on gene expression, nutrient allocation, and feeding avidity. A greater understanding of the mechanisms underlying dehydration tolerance may provide insight into how environmental factors contribute to the behavior and distribution of thrips and their infectious agents, and may lead to novel control methods.

## Methods

### Maintenance of *Frankliniella occidentalis*

Thrips were acquired from North Carolina State University that originated from the island of Oahu, HI (Schneweis et al. 2017). Thrips were reared in glass mason jars (Ball, 16 oz. Wide Mouth pint) with a 9 cm filter paper used as a lid to allow air flow and prevent escape. Thrips were maintained on green bean pods, *Phaseolus vulgaris*, that had been disinfected in a 10% bleach solution for 30 minutes before feeding. Rearing jars were held in large plastic deli containers (Fabri-Kal PK32T, 32oz) with a mesh lid as a second layer of containment. Containers were kept in a temperature-controlled Percival incubator (I-30VL) at 24°C and 50-60% relative humidity with a 12:12 h light-dark photoperiod. Thrips use plant matter as their egg laying substrate so, in order to obtain synchronized age groups, green beans were removed from jars containing adult thrips and placed into a new jar until emergence to allow tracking of development. All thrips used in this experiment were 2-3 weeks old from initial emergence based on our tracking of development, meaning they had been in the adult stage for 3-5 days.

### Conditions to induce dehydration

Relative humidity (RH) of 0-10% RH was generated by calcium sulfate (CaSO_4_; 1.5×10^−2^% RH; Toolson, 1978), 95-100% RH by double-distilled water, and 73-77% RH by a saturated salt solution of sodium chloride (NaCl) (Winston and Bates 1960) in a 5L glass desiccator. Individual thrips were housed in a 1.5 mL Eppendorf tube with a small hole drilled in the top for airflow (Benoit et al. 2005). This air hole was covered using a small KimWipe square to prevent escape and allow airflow. The tubes were suspended in the desiccator using a perforated plastic stand to prevent the tubes from contacting the salt solution. Thrips were weighed (to 0.001 mg) using an electrobalance (SD±0.2 μg precision; CAHN 25; Ventron Co., Cerritos, CA, USA). The thrips were removed from the tubes they were placed in using an aspirator and placed into a small aluminum cup (mass <20mg). As thrips are highly mobile, the aluminum cup was used as a means to contain them on the weighing pan long enough to determine their mass. This aluminum cup was weighed before every measurement and this mass was subtracted from the following measurement to determine the thrips’ mass. Anesthetics were not used and thrips were permitted to freely walk about in the aluminum cup on the weighing pan of the balance during mass determination. Prior to weighing, thrips were held at 100% RH and 24 °C, with no food, for 1 hour to minimize the effects of digestion, reproduction, and excretion on mass changes (Arlian and Ekstrand 1975).

### Assessment of water balance characteristics

As defined by Wharton’s (1985) general water balance equation (Eq. 1):

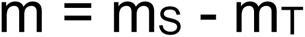

Total internal body water (m) is influenced by water moving out (transpiration, m_T_) and water moving in (sorption, m_S_). Negative mass changes indicate that more water is being lost than gained, or m_T_ > m_S_. Inversely, m_T_ < m_S_ indicates that more water is being gained than loss, resulting in a net gain in water mass. Therefore, m_T_ = m_S_ indicates that water balance has been achieved and there is no fluctuation in water mass.

The value of m, wet mass, was calculated by subtracting initial fresh mass (f) by the dry mass (d). To obtain the dry mass (d), thrips were weighed and frozen in a frost-free freezer (−20°C) to kill them and then placed in a drying oven at 0% RH and 50°C for 1 week and weighed. To ensure complete dryness, thrips were placed back into the drying oven and weighed 1 day apart until no mass change was observed. Percentage body water content was determined by the equation (Wharton 1985) (Eq.2):

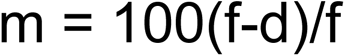

At 0% RH, there is no water available for sorption, meaning that m_S_= 0, thus any change in mass is due solely to the value of m_T_ (Eq. 1). Transpiration rate was assessed by weighing (fresh mass, f) and placing thrips in 0% RH and 24°C and weighing again after 4 hours (final mass, f_f_). The transpiration rate was determined to be the percentage loss per hour over this period (Eq. 3):

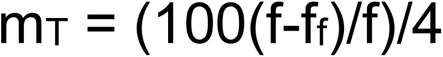

To assess dehydration-induced morbidity, thrips were placed into a desiccator held at 0% RH as described before and checked every 2 hours until all individuals had died. During assessment time, the entire perforated stand holding the tubes was removed to ensure all individuals spent the same amount of time outside of experimental conditions and that the relative humidity of the chamber was not adversely affected by being repeatedly opened and closed.

### RNA extraction of dehydrated thrips

Samples were collected and homogenized in large groups (N=60 males and N=20 females, three biological replicates per treatment) to reduce individual variation and increase RNA yield. Adult males and females were removed from the colony and placed into separate jars and kept under 75% RH and 24°C for 12 and 24 hours respectively. Different total times were used to account for the disparity in survival potential between the sexes as a means to observe the molecular interactions at a similar level of dehydration. Controls were placed at 100% RH for the same period of time. After treatment, twelve samples (three each of females and males with and without dehydration) were frozen at -80°C for at least 30 minutes. Individuals were separated out into tubes for extraction. Thrips were homogenized using a BeadBlaster 24 microtube homogenizer (Benchmark Scientific, Edison, NJ, USA). Total RNA was extracted using the RNAqueous Total RNA Isolation Kit (Thermo Fisher Scientific). RNA was treated with DNase I and then cleaned and concentrated using the GeneJET RNA Cleanup and Concentration Micro Kit (Thermo Fisher Scientific). Prior to sequencing, RNA concentration and quality were examined with a Nanodrop 2000 (Thermo Fisher Scientific) and Bioanalyzer (Agilent).

Poly(A) libraries were prepared by the DNA Sequencing and Genotyping Core at Cincinnati Children’s Hospital Medical Center. RNA was quantified using a Qubit 3.0 Fluorometer (Life Technologies). Total RNA (150–300 ng) was poly(A) selected for reverse transcription using a TruSeq Stranded mRNA Library Preparation Kit (Illumina). Multiplexing was conducted by ligating an 8-base molecular barcode to sequences before 15 cycles of PCR amplification and HiSeq 2500 (Illumina) Rapid Mode library sequencing. Sequencing resulted in 40–60 million paired end strand specific reads per sample. Reads are deposited to the NCBI under the following Bioproject PRJNA852191. Quality control of the raw sequencing reads yielded 539,522,498 reads, of which 57% mapped for males and 70% successfully mapped for females to the *F. occidentalis* reference genome per sample. 90% of the sequences had a Phred score over 35 and an average length of 93 base pairs.

### Bioinformatic analysis of transcriptional changes during thrips dehydration

Raw Illumina reads were processed through two independent pipelines for comparison (Davis et al. 2021, Pathak et al 2022). For the first pipeline, reads were trimmed for adapters in CLC Genomics Workbench version 12.0 (QIAGEN). The trimmed, paired-end reads were mapped to the official gene set version 1.1 of the *F. occidentalis* genome (Genbank assembly GCA_000697945.4, Focc_3) (Rotenberg et al. 2020). The mapped reads were then combined and normalized in CLC Genomics Workbench to generate normalized expression values as the number of transcripts per kilobase per million (TPM) as a proxy for expression. For the second pipeline, raw reads were uploaded to Galaxy (Afgan et al 2018), trimmed for adapters using Trimmomatic (version 0.38.0), and then mapped to OGSv1.1 and quantified using Salmon (version 1.5.1) to generate normalized expression values as TPM. Mapped read counts were summarized and combined using tximport (version 0.1).

Assessment of differential expression was conducted on the mapped reads of the 2 previously mentioned pipelines using 2 methods for comparison: the EDGE method with default settings in CLC Genomics Workbench 12 (QIAGEN) and EdgeR (version 3.34.0), for the CLC and Salmon pipelines respectively. False discovery rate (FDR) corrected p-values less than 0.05 were considered significant. Results of both pipelines were compared and only transcripts that were significantly differentially expressed in both were considered for further analysis. Pearson correlation coefficient was calculated to establish the overlapping differentially expressed transcripts from each pipeline.

### Gene ontology analysis of transcript change during thrips dehydration

Nucleotide sequences of all transcripts were subject to Blastx against the NCBI non-redundant protein database using the CloudBlast resource in Blast2GO v6.0.3 (BioBam) (Conesa et al. 2005, Conesa et al. 2008, Gotz et al. 2008). A Fisher’s Exact test was utilized to determine if our GO terms were occurring more often than would be expected within the set and was carried out using the native Fisher’s Exact test function within Blast2GO to identify differential enrichment of functional annotations with an FDR of <0.05.

### Nutrient reserve assays of dehydrated thrips

The effect of dehydration on nutrient reserves was assessed with assays for proteins, lipids, and glycogen based on methods developed for arthropod systems with low biological materials (Rosendale et al. 2019, Polak et al. 2017). Treatments were performed identically to those used in RNA-seq analysis for direct comparison to transcriptional shifts. Samples were collected and homogenized in large groups (N=60 males and N=20 females) and homogenized using a BeadBlaster 24 microtube homogenizer (Benchmark Scientific, Edison, NJ, USA). Soluble protein was measured using the Bradford method (BioRad; Hercules, CA, USA) with bovine serum albumin as the standard and absorbance read at 595 nm. Total lipid content was determined using vanillin reagent as previously described (van Handel 1985). Samples were homogenized in 2:1 chloroform:methanol. The solvent was evaporated between 80-90°C and then the remaining material was resuspended in H_2_SO_4_ and then continued to be heated at 80-90°C for 30 minutes. After cooling, 2 mL of Vanillin reagent (Sigma Aldrich) was added and the samples were read at 525 nm against a lipid standard of Crisco dissolved in chloroform (1 mg/mL). Glycogen content was measured using anthrone reagent (Sigma Aldrich) as previously described (van Handel 1985). Samples were homogenized in 1 mL of methanol and centrifuged at 5000 g for 5 minutes. The resulting pellet was resuspended in 1 mL of anthrone and then heated at 90°C for 7 minutes. Absorbance was read at 625 against a standard of glycogen dissolved in 25% ethanol (1 mg/mL). For all assays, 200 μl of the final homogenate was transferred to a 96-well plate and read using a Synergy H1 plate reader (BioTek, Winooski, VT, USA).

### Feeding Avidity Assay

The influence of dehydration on feeding behavior was assessed with the use of a dyed food resource. The contents of thrips’ gut can be seen through their exoskeleton; specifically, the food is dyed to increase contrast. Green beans were cut to a length of 3 cm and submerged in methylene blue (10 mg/mL) in a glass test tube for 2 days. The beans were removed from the tube and allowed to air dry for at least 1 hour before being presented to thrips. The green beans were distinctly blue in color, but touch did not transfer the dye. This indicates that only feeding can cause the blue coloration of thrips. Ten thrips of the same sex were randomly selected from the colony and placed into a small Mason jar (Ball) for treatment. Controls were placed in 100% RH and the dehydrated treatment groups were placed in 0% RH. Males were kept under treatment for 1.5 hours and females were kept in treatment for 3 hours to account for the differences in mortality and rates at which they lose water relative to their size. After the allotted treatment time passed, the thrips were provided with the dyed green bean. After 1 hour, the jars were frozen at -20°C in a frost-free freezer to kill the thrips and prevent further feeding. Thrips were individually assessed under a dissecting scope for blue coloration in the digestive system. Thrips were considered to have fed if any blue coloration was detected after this period. Feeding avidity was defined as the proportion of individuals that fed during a given time interval (League et al. 2021).

### Water balance and nutritional dynamics during infection

Infected thrips samples were provided from North Carolina State University. Briefly, the insect colony of *Frankliniella occidentalis*, originating from Oahu, HI, was maintained on the commercially purchased green bean pods (Phaseolus vulgaris) at 22°C (±2°C), 16L:8D photoperiod as described previously (Han & Rotenberg, 2021). Tomato spotted wilt virus (isolate TSWV-MT2) was maintained on *Emilia sonchifolia* by alternate transmissions between inoculation by viruliferous adult thrips and mechanical inoculation using thrips-inoculated leaf tissues. The systematically infected, symptomatic *E. sonchifolia* leaves were collected after 12 days post mechanical inoculation and used as virus source of inoculum for virus acquisition by larval thrips.

Cohorts of young, age-synchronized first instar larvae (L1s) were obtained from female oviposition chambers to generate TSWV-infected and non-infected thrips as described previously (Han and Rotenberg, 2021). Briefly, females were allowed a 24-hour oviposition period on healthy green bean pods in sealed colony cups. Females were removed to leave impregnated green beans to incubate for three days to allow for the first flush of larvae, which were brushed off (removed) to allow beans to incubate for an additional 16 hours to generate age-synchronized L1s. Half of the L1 cohort was exposed to symptomatic, TSWV-infected *E. sonchifolia* leaves, and the other half exposed to virus-free *E. sonchifolia* leaves for 24 hours at 25 °C. This published method reproducibly generates larval cohorts (second instar larvae, L2) with TSWV infection rates of no less than 90%, i.e., nine out of 10 L2s are virus-positive as determined by real-time quantitative reverse-transcription PCR (qRT-PCR) (Han and Rotenberg, 2021). The larvae from these two treatments were then transferred to the green bean pods in two separate containers. Thrips samples were shipped overnight to the University of Cincinnati for further analysis. After shipments were received, thrips were reared to adulthood as previously described. In infected thrips, we measured water loss rates and survival at 0% RH, water mass, feeding avidity, and glycogen reserves to assess the interactions between TSWV infection and thrips’ ability to maintain their water balance. These methods were conducted as described above for the dehydration experiment.

### Statistics

Replicates throughout all experiments are biologically distinct, individual samples. Sample sizes are listed in the appropriate figure legend. Statistical significance between treatments is noted within the results section or in the figure legend. Statistical tests are listed within the respective figure legends. Data processing was conducted in Microsoft Excel (v.2201) and R (v.4.0.4) (R Core Team, 2017) using RStudio (v1.4.1106) (RStudio Team, 2015). Statistics and graphical representations of data were performed in R using RStudio. Packages utilized include ggplot2, plyr, dplyr, AER, ecotox, Rmisc, RColorBrewer, emmeans, gam, mgcv, multcomp, pheatmap, and viridis. GO pie charts were generated utilizing CirGO in Python (v.3.9) (Kuznetsova et al. 2019).

## Results

### Water Mass (m) differences between males and females

Critical to forming a baseline of water balance dynamics in thrips, overall water content was first assessed. Average fresh mass (f) of males was 14.53 μg (± 1.88), and 42.07 μg (± 3.47) for females (ANOVA; P = 2.2 e-16). Dry mass (d) was 4.33 μg (± 0.72) and 14.33 μg (± 1.91) for males and females, respectively (ANOVA; P 2.2e-16). The overall percent water content was 69.82% (± 6.16) for males, and 66.01% (± 2.49) for females (ANOVA; P = 0.00087). Overall, while females are larger in overall size, males maintain a significantly larger overall percentage of their mass as water (Table 1).

**Table 1:** Comparison of water balance characteristics of adult male and female *F. occidentalis* (n=30). Statistics were performed using a one-way ANOVA and all values were significantly different between the sexes (P < 0.05).

| Characteristic | Male | Female |
| --- | --- | --- |
| Initial mass (µg) | 14.53 ± 1.88 | 42.07 ± 3.47 |
| Dry mass (µg) | 4.33 ± 0.72 | 14.33 ± 1.91 |
| Water mass (µg) | 10.2 ± 1.89 | 27.73 ± 2.05 |
| Body water (%) | 69.82 ± 6.16 | 66.01 ± 2.49 |
| Water loss rate (%/h) | 8.91 ± 3.04 | 4.39 ± 1.87 |
| Survivorship LT50<br>(Hours) | 9.81 ± 1.04 | 19.02 ± 1.34 |

### Water Loss (m_T_) and survival reveal sexually dimorphic response to dehydration

Transpiration rate was determined in order to assess thrips ability to retain water and resist desiccation. After a 4 hour dehydration bout where m_S_= 0, males lost, on average, 6.08 μg (±2.16) of mass. Females lost 8.27 μg (±3.24) in the same amount of time. Based on the average initial size for this experiment, where females weighed 48.43 μg (±8.19) and males weighed 17.16 μg (±2.56), females have approximately 2.8 times more mass than males do (ANOVA; P = 2.2e-16). Accounting for their total mass when looking at water mass, males lose 35.64% (±12.15) of their total mass, equating to a percentage loss of 8.911%/h (±3.04). Females under the same conditions lost 17.58% (±7.48) of their total mass, or 4.39%/h (±1.87). Thus, males lose a far greater amount of water (ANOVA; P = 4.86e-11) under desiccating conditions than females do relative to their body size and available water pool (Fig. 1A). General mortality under desiccating conditions was assessed to determine how well thrips tolerate dehydration over time. Kept under extreme desiccating conditions where m_S_= 0, females performed significantly better than the males (t-statistics; P = 3.56e-14). The LT50 for males was 9.81 h ± 1.04 and 19.02 h ± 1.34 for females, significantly in favor of the females as they survive twice as long on average (Fig. 1B).

**Fig. 1:**
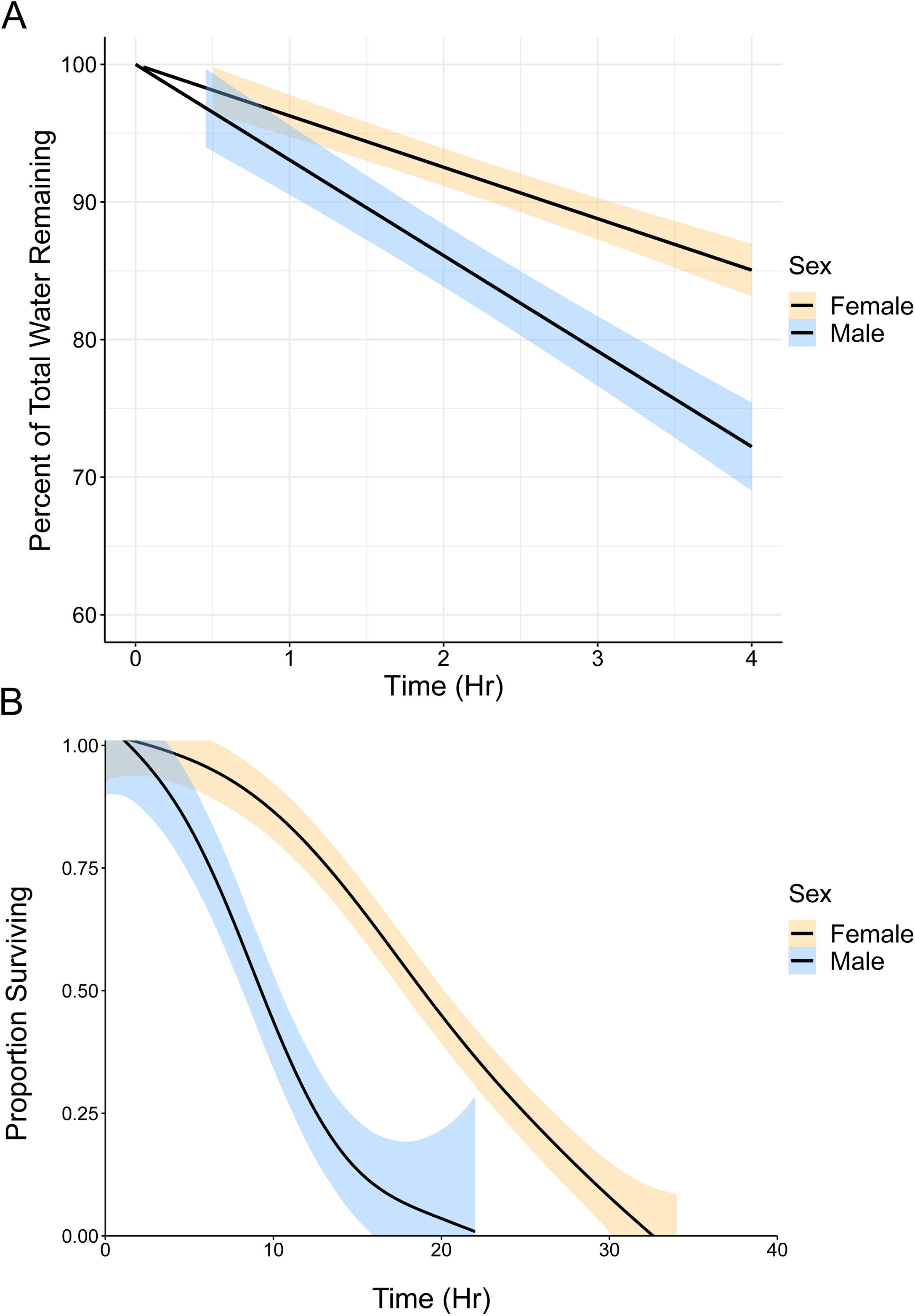
Water loss and survival rate in adult male and female western flower thrips (*Frankliniella occidentalis*) reveal sexually dimoprhic differences in dehydration tolerance. A) Percentage of water mass lost by living adult male and female *F. occidentalis* at 0% RH and 24 °C. Water loss rate is derived from the slope of the lines depicted on the plot. Values depicted are a mean of each treatment with 38 male and 48 female individuals and statistics were conducted using a one-way ANOVA (P < 0.05). B) Proportion of adult male and female thrips that survived 0% RH over time. Values are a mean of each treatment with 5 groups of 15 individuals; error is depicted utilizing a generalized additive model. Values were significantly different from each other (T-statistics, P < 0.05).

### RNA-seq analyses reveals difference in sugar metabolism

RNA-seq was used to elucidate the molecular underpinnings of dehydration tolerance in adult male and female thrips. Clustering the samples using principal component analysis (PCA), the greatest source of variation in the data set is attributed to the difference between males and females with PC1 explaining 63.4% of the variance in the dataset (Fig. 2A). The CLC pipeline yielded 1581 and 480 differentially expressed transcripts for dehydrated males and dehydrated females when compared to their controls, respectively (Supp. Table 1). The Salmon pipeline yielded 4007 differentially expressed transcripts for dehydrated males and 1466 for dehydrated females (Supp. Table 2). Comparing these pipelines, 1387 transcripts remained for dehydrated males and 436 transcripts remained for dehydrated females (Fig. 2B, Supp. Table 3). Between the males and females, there were 175 transcripts in common, both up and down regulated (Fig. 2C). A 0.823 and 0.832 Pearson correlation coefficient for males and females, respectively, indicated that the pipelines significantly overlapped (Fig. 2D). In order to determine the putative function of these transcripts, the predicted gene set (Rotenberg et al. 2020) was subjected to a blastx analysis against the NCBI nr protein database and GO categories were assigned using Blast2Go (Conesa et al. 2005). BLAST to nr yielded all but 137 of the 16717 transcripts with a significant BLAST hit. The Fisher’s Exact test was utilized to determine whether or not there is an overrepresentation of gene products that perform similar functions within a given list of transcripts. This was performed by inputting curated lists of differentially expressed transcripts to compare to the entire thrips predicted gene set (Rotenberg et al. 2020).

**Fig. 2:**
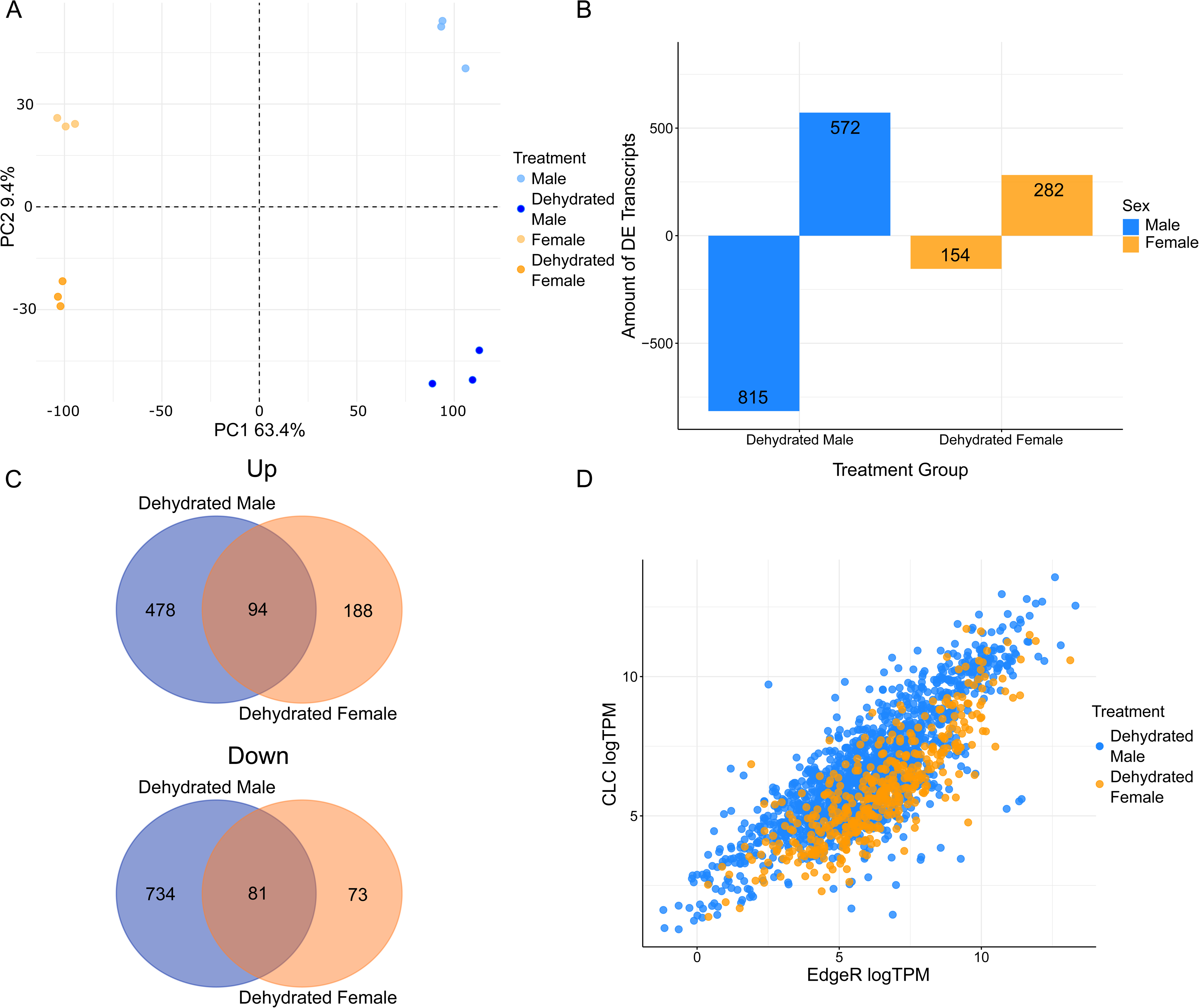
Sexually dimorphic differences in male and female differential expression. A) Principal component of dehydrated and control thrips RNA-seq samples. Each point represents one thrips homogenate (20 females or 60 males per homogenate). The directionality of the separation was observed in PC1 and PC2, accounting for 63.4% and 9.4% of observed variance, respectively. The separation of the points into distinct clusters indicates that the treatments were significantly different from one another. B) Total count of differentially expressed transcripts. Dehydrated males displayed a total of 572 upregulated and 815 downregulated transcripts and dehydrated females displayed a total of 282 upregulated and 154 downregulated transcripts. C) Overlap of differentially expressed transcripts in dehydrated male and female thrips. Dehydrated male and female samples were compared to their respective controls to determine the list of differentially expressed transcripts. D) Comparative distribution of differentially expressed transcripts logTPM across both pipelines in dehydrated males and females. Full list of differentially expressed transcripts in each pipeline are available in Supplemental Table 1 and 2. Full list of transcripts that appear in both pipelines are available in Supplemental Table 3.

When compiling all of the transcripts that were responsive across both males and females, there were 193 GO terms that were overrepresented across biological processes, molecular functions, and cellular components when compared to the entire predicted gene set (Fig. 3A). In general, there is a shared increase in carbohydrate metabolic factors during dehydration and a decrease in processes associated with generation and breakdown of proteins for males and females. As previously described, males had a larger amount of differentially expressed transcripts over females and the overlap between the sexes was somewhat small (Fig. 2B). When looking more specifically at the upregulated expression profiles of males and females, males exhibited more overrepresented GO terms than females with 42 terms for males versus 15 for females (Fig. 3B). For downregulated transcripts there is, again, a greater proportion seen in males than females. Males exhibited a sum of 155 overrepresented terms compared to the females’ 32 terms.

**Fig. 3:**
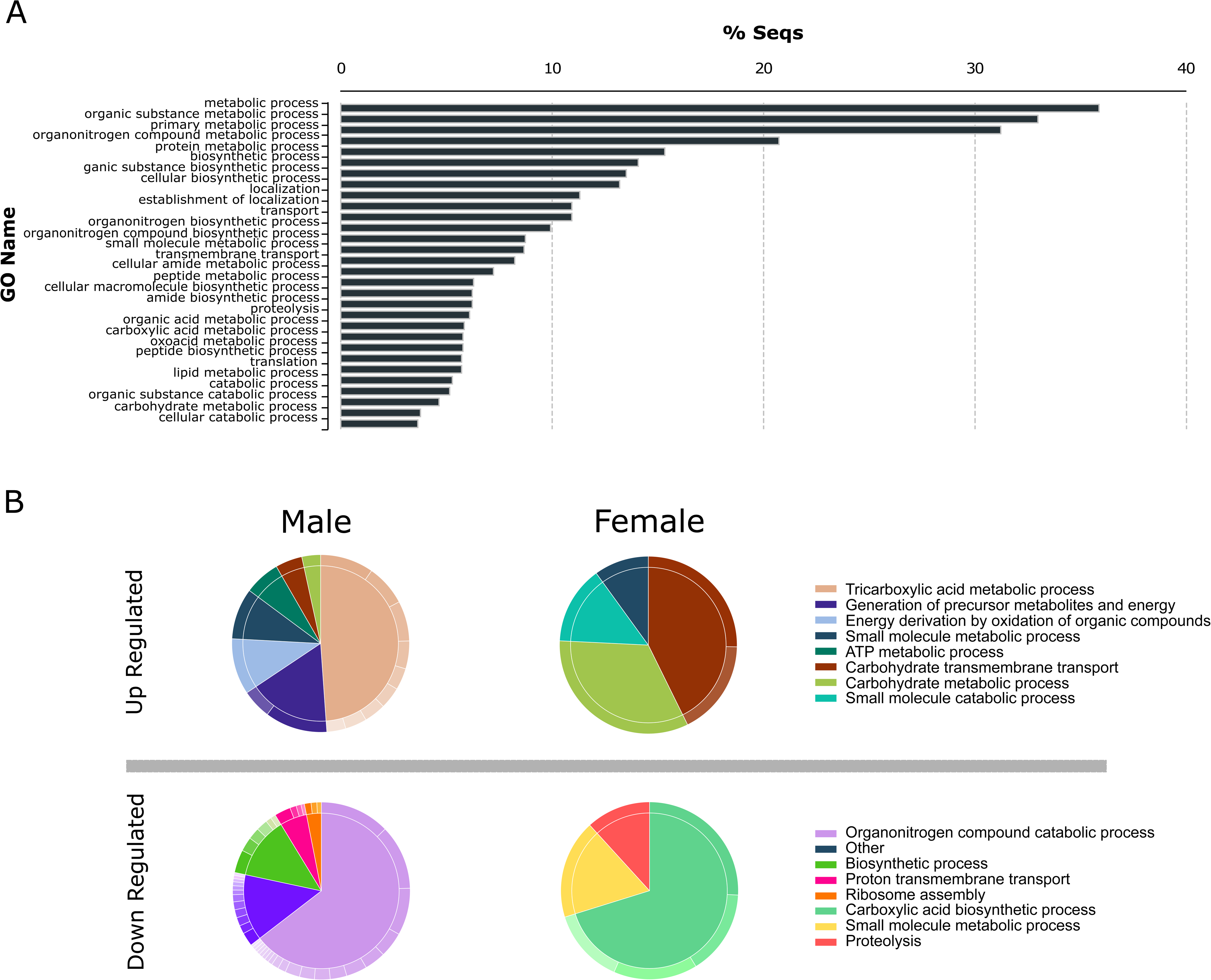
GO terms associated with transcriptional changes during dehydration in *F. occidentalis*. A) GO terms for the most abundant terms enriched among all of the 1823 differentially expressed transcripts. B) GO analysis of the biological functions of the significantly up and downregulated transcripts reveals a unique suite of functions in response to dehydration in males and females. Each significantly enriched group of GO terms is displayed as its most general categorization. The smaller slices of the pie on each group represent a more specific group of terms that are within that category of GO terms, which are available in Supplemental Table 4.

### Dehydration induced shifts in glycogen reserves

There was a significant enrichment for carbohydrate transport and metabolism in both males and females in response to desiccation based on the RNA-seq studies (Fig. 3). When quantifying the relative glycogen stores in adult thrips, there was a notable decrease after being subjected to desiccating conditions (Fig. 4A). Relative glycogen content in females was reduced by nearly 50% when exposed to dehydrating conditions (ANOVA; P = 0.0302). Males also experienced a reduction in glycogen content of nearly 30% when dehydrated, although not significantly reduced from the control (ANOVA; P = 0.359). When comparing the relative protein and lipid contents, the only other notable difference was a decrease in relative protein content found in dehydrated females, although not significant compared to control females (ANOVA; P = 0.162) (Fig. 4A). This difference was not found in males.

**Fig. 4:**
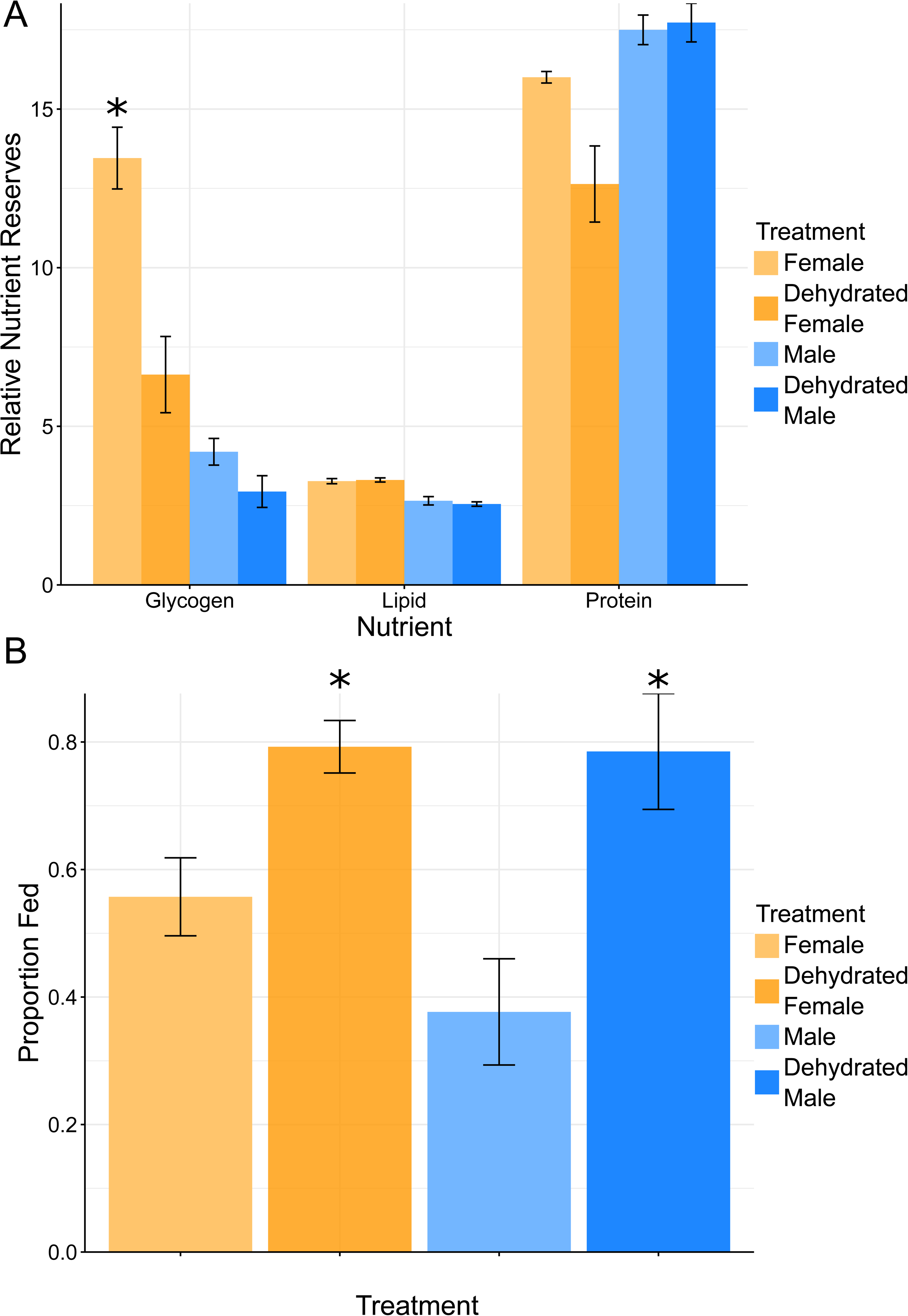
Dehydration induces a reduction in glycogen content and increases thrips feeding. A) Relative glycogen, lipid, and protein content in control and dehydrated males and females. Values are a mean ± SE of 3 samples each (each sample containing 20 females or 60 males). Statistics were performed using a one-way ANOVA. B) Feeding avidity of control and dehydrated males and females. Values are a mean ± SE of 3 trials each (each trial containing 10 individuals). Asterisks indicate values that were significantly different from controls. Statistics were performed using a one-way ANOVA (*P < 0.05).

### Feeding avidity increases after a bout of dehydration

Feeding avidity was assessed in order to gauge the influence of dehydration on thrips feeding behavior. When given 1 hour to feed after being under the stress of dehydration, 89.53% more males fed compared to those under control conditions (ANOVA; P = 0.0362). For dehydrated females, 42.2% more fed than females under control conditions (ANOVA; P = 0.0477). Thus, we can conclude that dehydration has a significant influence on the probability that a given individual will feed within 1 hour regardless of sex; however, a stronger response is seen in male individuals (Fig. 4B).

### Water balance and nutrition dynamics during infection

To determine the influence of TSWV infection on water balance characteristics, we repeated the experiments for overall water mass, water loss, survival, glycogen reserves, and feeding avidity in infected adults. When infected with TSWV, there was no significant difference between water mass ratios between infected and uninfected adult males (ANOVA; P = 0.981) or females (ANOVA; P = 0.272) (Table 2). Infected males had an average fresh mass of 13.73 μg (±2.43) and infected females had a fresh mass of 44.27 μg (±4.18). The dry masses were 4.33 μg (±1.05) and 15.2 μg (±1.37) for infected males and females, respectively. The overall body water percentage was thus 68.21% (± 6.4) for infected males and 65.55% (±2.54) for infected females. In terms of water loss, males lost 21.71% (±7.69) and females lost 14.83% (±8.59). Thus, each hour, infected males lost 5.43%/h (±1.92) and infected females lost 3.71%/h (±2.15) (Fig. 5A). Infected males (ANOVA; P = 1.25e-5), but not females (ANOVA; P = 0.1197), lost water less quickly than when uninfected. For survival, there is no significant difference between infected and uninfected males (T-statistics; P = 0.363) or females (T-statistics; P = 0.539), although there is a slight trend towards infected individuals being less hardy, as the LT50s were 7.46 h ± 1.30 and 15.86 h ± 1.83 for infected males and infected females, respectively (Fig. 5B).

**Fig. 5:**
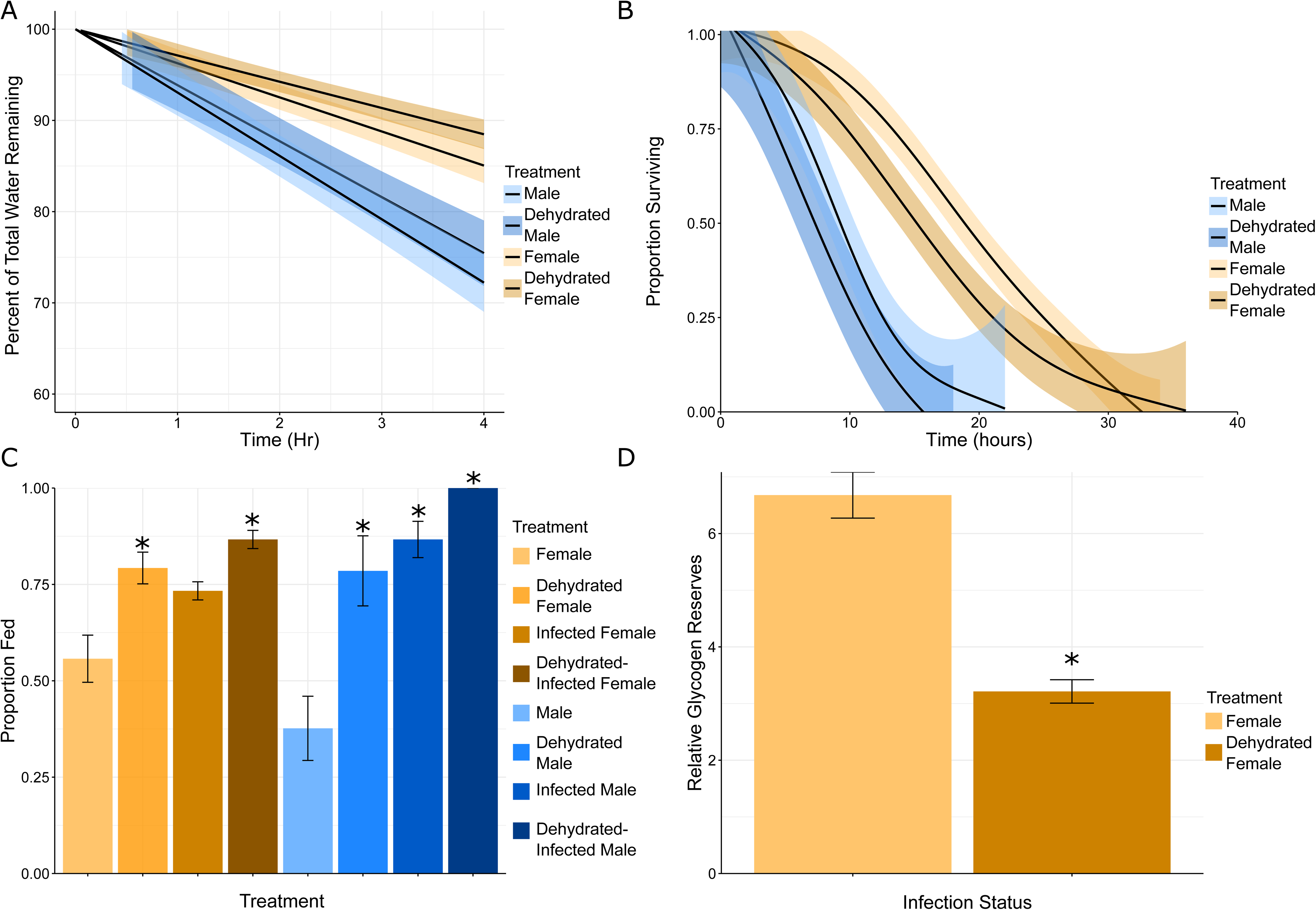
Infection induced effects on dehydration tolerance and nutrition dynamics. A) Percentage of water mass lost by infected male and female F. occidentalis compared to uninfected at 0% RH and 24 °C. Water loss rate is derived from the slope of the lines depicted on the plot. Values depicted are a mean of each treatment with 38 male and 48 female uninfected individuals and 15 male and 39 female infected individuals. Statistics were conducted using a one-way ANOVA. Males and infected males are significantly different from one another (P < 0.05). B) Proportion of infected and uninfected adult male and female thrips that survived 0% RH over time. Values are a mean of each treatment with 5 groups of 15 individuals for uninfected 2 groups of 15 for infected males, and 3 groups of 15 for infected females; error is depicted utilizing a generalized additive model and statistical analysis were conducted using T-statistics. C) Feeding avidity of control, dehydrated, infected, and dehydrated and infected males and females. Values are a mean ± SE of 3 trials each (each trial containing 10 individuals). Statistical analysis was conducted with a two-way ANOVA using a GLM that included dehydration and infection status as fixed effects and their interaction term followed by Tukey’s HSD post-hoc analysis. D) Relative glycogen of control and dehydrated infected females. Values are a mean ± SE of 3 samples each (each sample containing 10 females). Statistics conducted using a one-way ANOVA. Asterisks indicate values that were significantly different from controls (*P < 0.05).

**Table 2:**
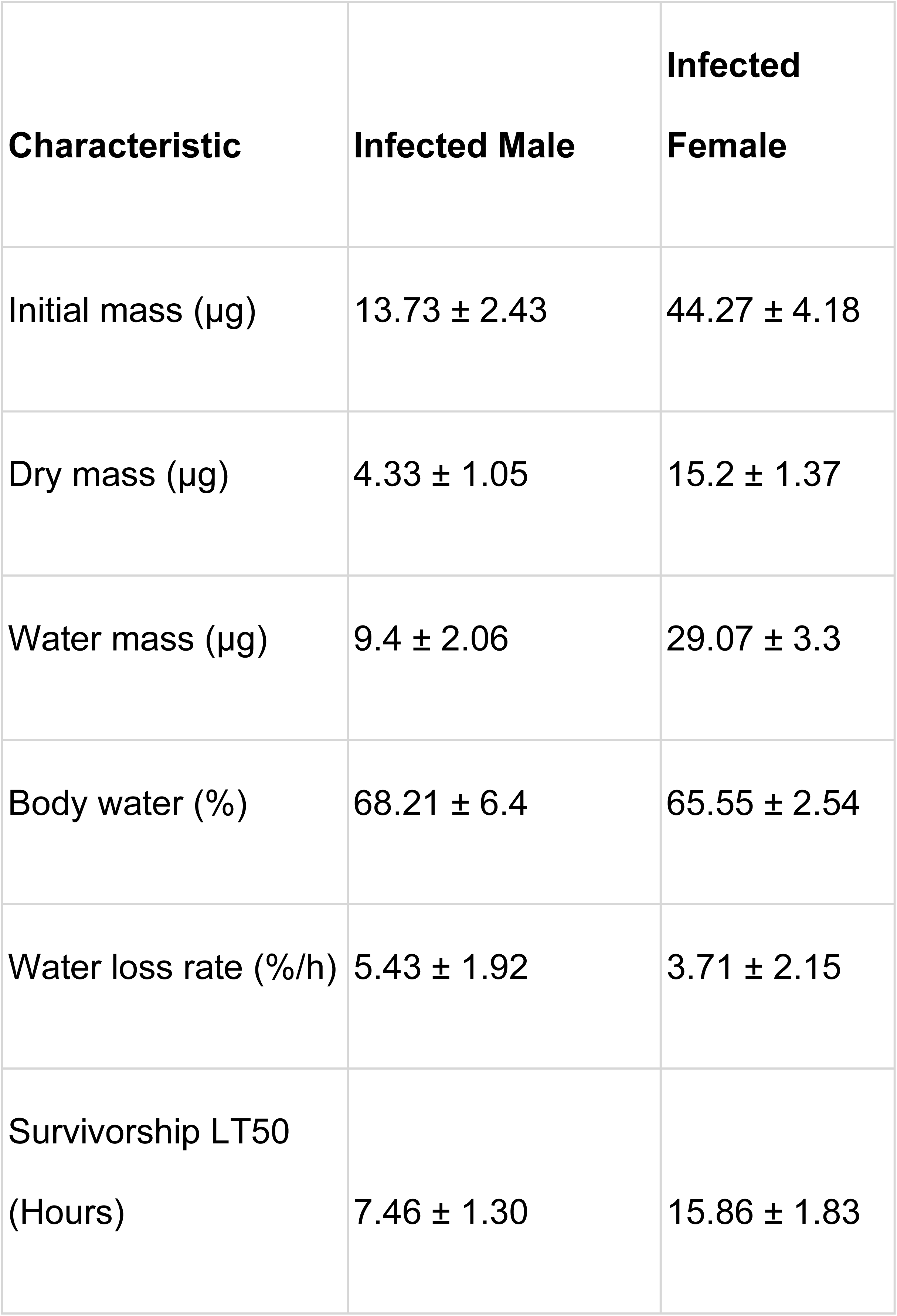
Comparison of water balance characteristics of infected adult male and female *F. occidentalis* (n=15). Statistics were performed using a one-way ANOVA and all values were significantly different between the sexes (P < 0.05).

The influence of TSWV infection was also assessed for feeding avidity and glycogen reserves. When comparing feeding avidity, in infected males 138.9% more fed (Tukey; P = 0.0151) and when infected males were also dehydrated, 165.5% fed more than the male control (Tukey; P = 0.0038). When females are infected, 31.6% more fed (Tukey; P = 0.1416) and when both infected and dehydrated, 55.5% more fed (Tukey; P = 0.0126). Despite trending higher, the influence of both dehydration and infection is not significantly different than either dehydration or infection alone for either males (Tukey; Infection P = 0.748, Dehydration P = 0.417) or females (Tukey; Infection P = 0.407, Dehydration P = 0.798), indicating that the influence of these factors together could not be determined to be additive effects to feeding in this experimental design (Fig. 5C). When measuring the relative glycogen reserves compared to uninfected controls, infected females displayed an average reduction of 51.84% (ANOVA; P = 0.0146). Thus, similarly to dehydration, infection has an adverse effect on the glycogen stores for females (Fig. 5D). These results indicate that both dehydration and infection are likely to increase feeding, potentially through similar mechanisms since glycogen reserves decreased significantly in both treatments, at least for females.

## Discussion

Our experiments established the basic water balance characteristics of adult thrips of both sexes, highlighting a distinct sexual dimorphism in overall size and percentage water mass. Visually, females are larger than the males at nearly three times the mass of males on average. Males maintain a higher proportion of their overall mass as water when compared to females. In terms of water loss, males lose water more quickly relative to their overall mass. Ability to survive under desiccating conditions heavily favors the females due to the males’ increased water loss. RNA-seq analysis of expression and analysis of nutrient reserves of thrips’ bodies provided supportive, complementary evidence for the hypothesis that both sexes of *F. occidentalis* exhibit enhanced carbohydrate metabolism from glycogen reserves under water stress in order to maintain water balance. Following dehydration, both males and females exhibited an increased likelihood to feed. The addition of TSWV-infection status as another factor in the dehydration experiment modulated water loss in males. Infected males, but not females, exhibited a significantly reduced water loss rate. Survival under desiccating conditions was not significantly affected for either sex. It is possible that with a greater sample size this effect would be statistically significant. In infected females, glycogen was reduced in a similar fashion to dehydrated females. When infected, thrips had an increased propensity to feed and when both infected and dehydrated, they still exhibited an increased feeding avidity but not significantly greater than just dehydration or infection alone. While not statistically supported in the current study, the average proportion of thrips that fed when they received the combined treatments of virus infection and dehydration tended to be greater than the proportions that fed experiencing a single treatment. Increasing the number of insect replicates or modification of the experimental protocol might reveal an additive effect on feeding behavior. Overall, the results revealed that both dehydration and, for females, virus infection negatively affected glycogen reserves, which in turn, likely leads to increased feeding activity as an effort to replenish the glycogen and water stores.

Adult male and female thrips are visibily sexually dimorphic, and as such we predicted that difference in body size would be reflected by mass, but not necessarily by water content. The average female had nearly three times the total mass of the average male. Of their mass makeup, males had a greater proportion of their body mass composed of water (∼70%) than females (∼65%), which is within a similar range for some flies and mosquitoes (Arlian et al. 1975, Benoit et al. 2007, Gray and Bradley 2005). Thrips are diminutive in size, indicating that they likely experience significant loss of water through their cuticles due to their high surface area to volume ratio (Wharton 1985, Chown and Nicholson 2004). We predicted that both sexes of thrips would have a high transpiration rate and thus be defined as a hygric species (high water loss rates, usually over 2.0%/h), based on the classification system proposed by Hadley (1994). Indeed, both male and female thrips would be considered to be hygric because they have a relative water loss rate >2.0%/h (Hadley 1994). Compared to other hygric species, such as mosquitoes, western flower thrips lose a similar amount of water (Benoit et al. 2007). Water loss rates are an important indicator for assessing an insect’s suitability for any particular habitat (Wharton 1985, Hadley 1994). In contrast to hygric species, insects that live in arid regions and spend a prolonged amount of time off-host tend to be adapted to retain water better (Hadley 1994, Benoit and Denlinger 2010). The incredibly high transpiration rate displayed by thrips provides evidence that retaining water is not a strategy utilized by thrips in order to maintain water balance as they, instead, focus heavily on water uptake through feeding like other hygric species, such as mosquitoes (Hagan et al 2018). Their high transpiration rate is not surprising since insects that live directly on their food source, whether plant or animal, tend to have high water loss rates (Burgess 2022). Comparison of males and females indicated that the males lose water significantly faster under the same conditions, most likely due to the size advantage in favor of females. This difference in water content across sexes is not particularly uncommon, as a similar trend is seen in both *Drosophila* and mosquitoes where the females are also larger than the males (Gibbs and Matzkin 2001, Eckstrand and Richardson 1980, Benoit et al. 2007). Due to the previously aforementioned high transpiration rates, we predicted that the overall ability to tolerate extreme desiccation would be poor, and that females would survive significantly longer than males due to their lower relative loss of water. Indeed, mortality was significantly higher in males, with the LT50 being less than half of what was displayed by females. Since males are smaller than females, they not only have a smaller surface area to volume ratio, but they also have less overall water to lose and smaller reserves of carbohydrates to metabolize than females do (Chown and Nicholson 2004), resulting in a lessened ability to sustain water balance under severe dehydration pressure by breaking down glycogen into molecular water (Loveridge and Bursell 1975). It is not uncommon for insects that have a lower overall water content, in this case the females, to be more tolerant of desiccation (Hadley 1994). This is likely because increased dry mass also increases the size of the insect, which has been observed by the increased desiccation tolerance seen in diapausing mosquitoes (Benoit et al. 2007).

Overall, the comparison of responses revealed that the defense against desiccation is highly sex-specific which can likely be attributed to the differences in overall desiccation tolerance, as females do not experience the same extreme pressure under desiccating conditions than males do. Given the many sexual dimorphisms already noted between males and females, through RNA-seq it is undoubtedly valuable to separate the male and female specific transcripts to see in what way males and females respond similarly and differently. As noted by PCA, the majority of variation across the sequencing samples is attributed to the difference between males and females, rather than the effect of dehydration. Males also experienced a greater degree of change when dehydrated. Despite this there was enrichment of Gene Ontology terms associated with carbohydrate metabolism for both males and females, which has been noted in other terrestrial arthropods (Hagan et al. 2018, Teets et al. 2012, Rosendale et al. 2016). Fluctuations in carbohydrate levels in response to dehydration have been documented in other insect systems to prevent additional water loss and reduce potential damage due to desiccation (Michaud et al. 2008, Teets et al. 2012, Rosendale et al. 2016). Carbohydrate metabolism could also possibly pertain to the breakdown of carbohydrates, particularly glycogen reserves, to form water, which is a strategy for maintaining water balance in other arthropods (Loveridge and Bursell 1975). Alternatively, carbohydrates have colligative properties and can be accumulated to act as osmoprotectants, helping to conserve water, preventing potentially damaging biochemical processes (Teets et al. 2012, Toxopeus et al. 2019). Males, but not females, displayed upregulation for terms related to energy conversion via TCA cycle and ATP metabolism, implying that this level of dehydration is particularly energetically costly for males. Females, on the other hand, displayed neither an upregulation nor downregulation of factors regarding energy production. The increase in males and lack of change in females indicates that there is likely no overall metabolic suppression like that which is commonly seen in other arthropod systems (Marron et al. 2003, Benoit et al. 2007). The response seen in males is more like that which is seen in ticks and spider beetles, which presumably increase their metabolic rate in order to produce metabolic water and defend against dehydration (Rosendale et al. 2016, Benoit et al. 2005). This is surprising because the lack of metabolic suppression is more commonly observed in arthropods with low water loss. Since thrips likely have nearly continuous access to plant food sources, metabolic generation of water could be a useful strategy to survive short periods of dehydration. Of particular interest, males also exhibited some upregulation of cuticle structural constituents in response to desiccation. Changes in cuticular structure and increased melanization have been noted in other arthropods as a method of increasing water retention (Benoit 2010, Rajpurohit et al. 2008, Parkash et al. 2008), highlighting a potential increased adaptive resistance to desiccation across multiple bouts of dehydration (Benoit et al. 2010). As thrips also display rapid cold hardening when given the chance to acclimate to low temperatures (McDonald et al. 1997), it is certainly within the realm of possibility that they are able to acclimate to drier conditions as well. Downregulated transcripts in males and females infer reduced proteolysis and fatty acid biosynthesis as common GO terms. These down regulated terms may indicate a shift in macromolecule priority, focusing on carbohydrates, rather than proteins or lipids, in order to resist desiccation in the short term. Additionally, the downregulation of proteolysis and fatty acid biosynthesis could hint at a potential state of quiescence while under dehydration stress in order to conserve water, although this behavior is often seen in insects with lower base water loss rates (Benoit et al. 2005, Benoit et al. 2007). Interestingly, proteolysis has been seen in other similar studies to be upregulated during dehydration stress as a means to presumably break down and replace damaged proteins (Rosendale et al. 2016). Additionally, a fairly common mechanism used by many insects to reduce water loss is the accumulation of cuticular hydrocarbons (Benoit et al. 2007, Gibbs et al. 1998). The lack of upregulated GO terms pertaining to proteolysis and cuticular hydrocarbons indicate that these are potentially not strategies utilized by thrips in order to resist water loss and damage due to desiccation. Importantly, the accumulation of cuticular hydrocarbons is more commonly associated with population differences between dehydration resistance or in individuals entering dormancy (Wang et al. 2022, Kellerman et al. 2018), thus rapid changes during dehydration are uncommon.

As highlighted by RNA-seq, there was an enrichment for transcripts that are putatively noted to be involved in carbohydrate transport and metabolism after being exposed to desiccating conditions. There was also a notable down regulation in factors related to lipid and protein metabolism and transport, indicating that processes involved with these macromolecules were being reduced in favor of focusing on carbohydrates. Thus, it was expected that, under the same pressure, we would be able to detect a reduction in glycogen stores in dehydrated thrips. Indeed, the glycogen reserves were significantly reduced in dehydrated females. For males, the glycogen was reduced by nearly 30%, but was not statistically significant. Potentially, increased sampling may lead to a statistically significant difference in glycogen in dehydrated males. In *Drosophila*, rates of lipid and protein metabolism were not altered during desiccation, but carbohydrate metabolism was several-fold higher (Marron et al. 2003). Additionally, *Drosophila* populations that were selected for dehydration resistance had a baseline increased store of glycogen (Djawdan et al. 1998, Marron et al. 2003). Interestingly, the protein content in dehydrated females, but not males, was also reduced. The lack of a change in males is not surprising because the majority of downregulated transcripts pertained to organonitrogen catabolism. Females, however, displayed a significant downregulation for proteolysis, making this decrease in protein content in dehydrated individuals somewhat surprising. It is important to note that this difference in protein quantity between males and females may be a result of experimental design; in our case, we wanted to attempt to create a similar level of dehydration for both the males and females to be observed through RNA-seq. Since males lose water much more quickly and survive, generally, half as long, halving the treatment time for males would potentially allow for insight into the transcriptome for males and females at a similar dehydration state. Given the length of time females spent under treatment, it is possible that, at some point, there is an increase in proteolysis or organonitrogen catabolism between the 12 hour and 24 hour experimental time points. Alternatively, it is possible that females simply respond differently than the males on a molecular level when they reach a similar dehydration state, given their apparent sexual dimorphism. Lipid levels were not altered in either males or females, similarly to other studies (Teets et al. 2012). As previously stated, the primary metabolic strategy for maintaining water balance in situations where water is scarce is likely the breakdown of glycogen as a source of water (Loveridge and Bursell 1975, Rosendale et al. 2017). Given their high relative transpiration rate, glycogen breakdown is likely the first resort in a short term bout of dehydration. For longer term dehydration, if thrips are, in fact, able to acclimate to better resist dehydration, we would expect to see a shift toward lipid metabolism as they are vital to increasing water retention properties (Gibbs et al. 1998).

Dehydration is a factor that can increase feeding behaviors in order to maintain water balance via increased water uptake (Hagan et al. 2018, McCluney 2017). Our prediction was that dehydration would significantly increase the proportion of individuals that had fed after 1 hour, regardless of sex. Dehydration did, in fact, increase the proportion of feeding individuals. In males, this effect was more pronounced, likely due to the tendency for males to probe food more often and less intensely than females (Ogada and Poehling 2015). For insects that feed to rehydrate, the strategy for surviving this lifestyle requires that they either feed often or are able to sustain long periods without any water uptake (Benoit et al. 2015). As we displayed, their high water loss rate indicates that thrips do not rely on their ability to conserve water and likely need to feed often. This implies that thrips feeding may increase significantly in the event of a drought in order to sustain their water balance; however, our experimental design only looked at the influence of dehydration in regards to thrips. In the field, the plants that thrips feed on are also exposed to the same desiccating conditions. Under these stressful conditions, plants may potentially become more nutritious or have reduced defenses to insect attack (Dale et al. 2017, Farooq et al. 2009), likely leading to a further increase in feeding. Further studies targeting plant-pest interactions should take this into account to help model and assess the potential increase in pest damage during moderate to severe drought. Humidity, or the lack thereof, is also responsible for other behavioral changes in many insect species: for example, many insects search for reprieve when humidity is low (McCluney 2012). Oftentimes, many species will retreat to the microhabitat with the highest relative humidity during the driest periods of the day (Kessler and Guerin 2008). Most likely, many species are able to detect humidity gradients, which is vital for protection against desiccation (Enjin et al. 2016). It is not definitively known whether or not thrips can detect relative humidity; however, they do possess the genes for the ionotropic receptors that are required for thermo and hydrosenstation in other insect species (Knecht et al. 2016, Rotenberg et al. 2020). Future researchers may be interested in determining thrips’ ability to detect shifts in relative humidity and choose more favorable conditions.

Since TSWV must be obtained during a larval stage to be spread (Wetering et al. 1996), the individuals in this experiment were inoculated with the virus early in life. Since viruses are known to be able to modulate host physiology in other insect species (Porras et al. 2020, Xu et al. 2016, Pusag et al. 2012), it was hypothesized that early onset infection could potentially influence adult thrips’ water balance characteristics in the long term as it persists through development. Additionally, TSWV infection has been implicated to down regulate proteins associated with cellular trafficking and movement of molecules throughout the organism in first instar larvae (Badillo-Vargas et al. 2012), which could potentially disrupt water balance through development. Overall, we predicted that infection would result in multiple viral modifications that ultimately led to an increased requirement for water uptake, as increased feeding probes have been reported previously in males (Stafford et al. 2011). Since glycogen was seen to be reduced in dehydrated female thrips, which also exhibited increased feeding avidity, we additionally predicted that there was a reduction in glycogen for infected thrips that possibly explained the increased feeding behavior. In summary, we predicted that infection would result in decreased overall water content, increased transpiration rate, decreased survival, increased feeding activity, and reduced glycogen content. There are a few studies that postulate that host-virus or host-parasite interactions may be affected by dehydration, importantly manifesting in behavioral differences that may increase disease spread (Hagan et al. 2018, Rosendale et al. 2016, Bezerra Da Silva et al. 2019). Other studies looking at similar host-virus interactions in insects reported that there are differences in thermal tolerance, preference, and performance at different temperatures, although these effects varied heavily across different host-virus systems (Porras et al. 2020, Xu et al. 2016, Pusag et al. 2012). The presence of these modifications indicate that similar modifications could be present for water balance characteristics. In particular, as cross tolerance between thermal and dehydration stress has been documented in numerous insect systems, it is entirely possible that the changes in thermal tolerance could simultaneously influence desiccation tolerance (Sinclair et al. 2013, Rosendale et al. 2016). The overall water content was not significantly different between infected adults and uninfected adults, indicating that TSWV does not likely modulate the way that thrips store water. This eliminates the potential that early onset infection significantly manipulates adult thrips water content through development; however, it is still possible that the water content could be altered during different life stages as thrips develop. When infected, the transpiration rate was unexpectedly lower than when uninfected, significantly so in males. This indicates that TSWV may be, in some way, increasing thrips’ water retention. Despite decreasing overall transpiration rate, there was no significant difference in the survivorship between infected and uninfected individuals. This hints that there is a tradeoff associated with viral infection whereby another unmeasured variable may limit their ability to survive desiccation. Previous transcriptomic analysis of the influence of TSWV through development indicated a large-scale down regulation in cuticle related proteins and factors (Schneweis et al 2017). This transcriptional shift was determined to be due to virus modulation in order to better disseminate from the midgut, but there was also postulation that this also likely influenced the outer cuticle as well. Transpiration through the cuticle is a main avenue for water loss which can be mitigated through modifications of the cuticle following exposure to bouts of dehydration (Benoit et al. 2007, Benoit 2010, Gibbs et al. 1998). Potentially, virus modulation may inadvertently be influencing the cuticle in such a way that thrips reap the benefit of increased resistance to desiccation, offering a possible explanation as to why there was an observed difference in water loss rates.

Similarly to the effect of dehydration, there was a notable reduction in glycogen content in infected females. This represents a strong pressure to feed more often to replenish glycogen stores, even outside of stressful conditions. Although only characterized in infected females, glycogen loss could potentially offer vital insight into the driving factors that contribute to the increased feeding probes previously seen in infected male thrips (Stafford et al. 2011). When looking at feeding avidity, infected males did, in fact, feed at a higher proportion than control individuals. Since both dehydration and infection lead to increased feeding by way of a reduction in glycogen stores, it is likely that this interaction is additive, although our experimental design could not parse out that difference. Minor dehydration in infected individuals may be additive; however, the more significant dehydration that we incurred increased feeding so much that we were unable to show differences. A previous RNA-seq study of TSWV infected thrips found a shift in expression of transcripts relating to carbohydrate metabolism in infected adults (Schneweis et al. 2017). This shift, although notably much smaller than in our dehydration treatment, may be attributed to the reduced glycogen content measured in infected individuals; however, this needs to be empirically determined. As viral infection is a constant pressure sustained by the host, it likely comes with a high energetic cost that drains the host’s energy stores (Arnold et al. 2013). A recent study of the *F. occidentalis* salivary glands when infected with TSWV reported an increase in proteins with functions relating to energy generation and protein turnover, providing evidence that infection with TSWV has a cost associated with it to thrips through damage to the salivary glands (Rajarapu et al. 2022).

## Conclusions

Despite their status as a globally invasive pest and significant disease vector, the physiological stress tolerances of thrips are understudied. Here, we characterized thrips’ dehydration tolerance as it is one of the most vital abiotic stressors that influence the abundance and distribution of insects in a given environment (Chown and Nicholson 2004). In addition to this, previous studies on mosquitoes revealed that dehydration increased feeding (Hagan et al. 2018). Thrips, due to their diminutive size, are particularly at risk for dehydration. Understanding how thrips resist desiccation, and the molecular factors at play could provide insight for predicting the spread of their range and provide novel control methods.

The work presented here provides baseline water balance characteristics and desiccation tolerance to further the understanding of thrips physiology, as well as the potential for insect host modulation by TSWV. This study focused primarily on the influence of single bouts of extreme dehydration; however, in reality, organisms contend with fluctuations of humidity and likely experience multiple bouts of mild desiccation (Benoit et al. 2010). As an invasive pest, understanding how thrips contend with dehydration long term is vital information to best understand their spread over time. For an insect as small as thrips, it is likely that they contend with fluctuations in humidity because they are able to effectively utilize microclimates to avoid uninhabitable conditions (Kessler and Guerin 2008). As such, future studies should focus on the ability of thrips to survive through sustained repeated bouts of dehydration. Additionally, other insects have displayed cross tolerance between thermal stressors and dehydration (Benoit et al. 2009). Since thermal stress and dehydration are some of the most important abiotic stressors that restrict an organism’s range, understanding the interactions between them would be invaluable (Chown and Nicholson 2004). Future studies should also aim to include infection dynamics to attempt to elucidate exactly how TSWV influences thrips. Infection has been noted to influence thrips, directly or indirectly, in several ways, such as their fecundity, development, population growth, survival, and feeding (Carter, 1939, Maris et al. 2004, Belliure et al. 2004, Shrestha et al. 2012, Ogada et al. 2012, Stafford et al. 2011). These results provide evidence that infection may increase thrips’ ability to retain water, but most certainly does not increase their hardiness under desiccating conditions without a food source. We found that infection reduces glycogen content which is most likely a main cause leading to the increased feeding seen in infected thrips. Due to this, a link between infection status and starvation may provide exciting insight into how this virus modulates host physiology. Ultimately, this creates a framework to allow future studies to evaluate thrips’ physiological stress limits and promote the inclusion of TSWV infection in such studies to increase the understanding of how thrips handle stress with and without the diseases they carry.

## Acknowledgements

We thank Alekhya Kondragunta and Hyojin Choi for assistance in data collection. Melissa Uhran, Oluwaseun Ajayi, A. Kondragunta, and H. Choi all assisted in animal husbandry. Christopher Holmes helped with R-script, Python-script, and figure construction. Funding was provided through the USDA (2019-67013-28495).

## Figures and tables

**Supplemental Table 1: EdgeR pipeline significant transcript list.** Full list of differentially expressed transcripts for both dehydrated males and females when compared to their respective controls.

**Supplemental Table 2: CLC pipeline significant transcript list.** Full list of differentially expressed transcripts for both dehydrated males and females when compared to their respective controls.

**Supplemental Table 3: Overlapped pipeline significant transcript list.** Full list of differentially expressed transcripts that appear in both Supplemental Table 1 and Supplemental Table 2. These transcripts are those that were used for further analysis.

**Supplemental Table 4: Full list of GO terms.** Entire list of Blast2GO output for All combined GO, Male Up, Male Down, Female Up, and Female Down.

